# An interaction between cancer progression and social environment in Drosophila

**DOI:** 10.1101/143560

**Authors:** Erika H. Dawson, Tiphaine P.M. Bailly, Julie Dos Santos, Céline Moreno, Maëlle Devilliers, Brigitte Maroni, Cédric Sueur, Andreu Casali, Beata Ujvari, Frederic Thomas, Jacques Montagne, Frederic Mery

**Affiliations:** Evolution, Génomes, Comportement & Ecologie, CNRS, IRD, Université.Paris-Sud, Université Paris-Saclay, 91198 Gif-sur-Yvette, France.; Institut for Integrative Biology of the Cell (I2BC), CNRS, Université Paris-Sud, CEA, UMR 9198, 91190 Gif-sur-Yvette, France; Département Ecologie, Physiologie et Ethologie, Centre National de la Recherche Scientifique, 67037 Strasbourg, France; Institut Pluridisciplinaire Hubert Curien, Université de Strasbourg, 67037 Strasbourg, France; Institute for Research in Biomedicine (IRB Barcelona), the Barcelona Institute of Science and Technology, Baldiri Reixac, 10, 08028 Barcelona, Spain; Centre for Integrative Ecology, School of Life and Environmental Sciences, Deakin University, Waurn Ponds,3216, Australia; CREEC, MIVEGEC, UMR IRD/CNRS/UM 5290, 34394 Montpellier, France

**Keywords:** Social interactions, group composition, cancer, Drosophila, disease ecology, behavioral ecology, parasite-host dynamics, evolutionary ecology of cancer

## Abstract

*This preprint has been reviewed and recommended by Peer Community In Evolutionary Biology (http://dx.doi.ors/10.24072/pci.evolbiol.100030)*

The influence of oncogenic phenomena on the ecology and evolution of animal species is fast becoming an important research topic. Similar to host-pathogen interactions, cancer negatively affects host fitness, which should lead to the selection of host control mechanisms, including behavioral traits that best minimize the proliferation of malignant cells. Social behavior is one such trait, which is suggested to influence cancer progression. While the ecological benefits of sociality in gregarious species are widely acknowledged, only limited data are available on the role of the social environment on cancer progression. Here, we exposed adult *Drosophila*, with colorectal-like tumors, to different social environments. First, we show that cancerous flies kept in complete isolation exhibit increased tumor progression. Yet, more surprisingly, we find that cancerous flies, kept in groups with other non-cancerous individuals, also develop tumors at a faster rate compared to those kept with other cancerous conspecifics, suggesting a strong impact of social group composition on cancer growth. Finally, we show that flies can discriminate between individuals at different stages of tumor growth and selectively choose their social environment accordingly. Control flies actively avoid flies with cancer but only at the later stages of tumor development, whereas cancerous flies display strong social interactions with cancerous flies in the early stages of tumor growth. Our study demonstrates the reciprocal links between cancer and social interactions, as well as highlighting how sociality impacts health and fitness in animals and its potential implications for disease ecology and ecosystem dynamics.

## Introduction

In gregarious species sociality not only offers important positive benefits associated with reducing predation risk (1) and increasing foraging efficiency (2), but also provides additional adaptive benefits by reducing overall metabolic demand (3), providing thermal advantages (4), decreasing stress responses (5) and increasing disease avoidance (6). It is therefore generally accepted that an individual’s social environment affects a large range of behavioral, psychosocial, and physiological pathways. Limited empirical evidence (mostly based on human studies) suggests that extreme social environments such as complete isolation or overcrowding of conspecifics in a group can potentially induce and accelerate pathological disorders. For example, in mammals, social isolation has been associated with faster progression of type 2 diabetes (7), cardiovascular or cerebrovascular disorders (8), and, notably, early and faster mammary cancer development (9, 10). Moreover, social overcrowding has been found to induce psychiatric and metabolic disorders (11). Few human studies have attempted to explore the role of social interactions on cancer progression, and the topic remains controversial. Adverse psycho-social factors, including traumatic life events, high levels of depressive symptoms, or low levels of social support, have been related to higher rates of, for example, breast and colon cancers (12, 13). However, these community based studies or meta-analyses often suffer from the complexity of inter-correlated factors. For example, low sample sizes, high risk behaviors associated with stress (e.g. smoking), and the heterogeneity and retrospective origins of these studies make it difficult to find a conclusive causal relationship between cancer progression and social conditions.

In addition to human studies, laboratory based experiments on gregarious species (e.g. Sprague-Dawley rats) have demonstrated an association between persistent social isolation and inflammatory responses linked to numerous disease processes, including cancer (9, 10, 14). Despite cancer (both transmissible and non-transmissible) being an emerging important factor influencing life history traits, even at early stages (15-18), little is known regarding the reciprocal links between the social environment and the development and progression of this illness. Increasing evidence demonstrates that oncogenic phenomena are extremely prevalent in host populations, and not just in post-reproductive individuals as previously believed (19). It is still largely unclear for both animals and humans how specific social group composition can directly affect tumor progression, and vice versa.

*Drosophila* has proven to be a powerful model system to address these issues. Social interactions are an important life history trait, particularly in female flies, who use social information to make fitness enhancing decisions (20-22). More importantly behavioral and physiological processes have been found to be influenced by the degree of social interaction while eliminating all other confounding variables. In *Drosophila*, social isolation leads to a reduced lifespan (23), increased aggression (24-26), reduced need for sleep (27, 28) and a decrease in the fiber number of the mushroom bodies in the integrative nervous center (29). Furthermore, tumor-like over-proliferation of tissues has been found to occur naturally in Drosophila (30, 31) and induced tumors have also been found to influence fitness traits in individuals (16, 18).

Here, to explore the reciprocal relationship between social environment and cancer progression, we made use of a colorectal-like tumor model (32). Thanks to genetic tools, tumors can be induced at a precise adult developmental stage and followed over fly lifespan. The tumors are generated by inducing clones in intestinal progenitor cells that are homozygous mutants for the two *Drosophila Apc (Adenomatous polyposis coli)* genes and that express an oncogenic form of the proto-oncogene Ras. Interestingly, loss-of-function of the APC tumor suppressor and expression of oncogenic Ras are critical steps to malignancy in the human colorectal track (33). First, we exposed tumor-bearing *Drosophila* females to various social environments for 21 days and measured tumor growth and social interactions. Subsequently, we tested control and cancerous flies for their social environment preferences predicting that cancer flies should presumably prefer the social environment which limits cancer progression.

## Results

### Biological model

Flies bearing heat shock-induced MARCM (Mosaic analysis with a repressible cell marker) clones (34) induced in 3-day old adult virgin females intestinal progenitor cells were used. The clones were mutant for both *Drosophila APC* genes, *Apc* and *Apc2*, and expressed the oncogenic form of Ras, Ras^V12^ and the GFP marker (Apc-Ras clones) (32). It has been shown that these compound Apc-Ras clones, but not clones either expressing Ras^V12^ or mutated for the *APC* genes, expand as aggressive intestinal tumor-like overgrowths that reproduce many hallmarks of human colorectal cancer (32). The number of GFP-positive gut cells was monitored with age every 7-days by flow cytometry from flies bearing either Apc-Ras or neutral clones, hereafter referred to as cancerous and control flies, respectively. As expected, a clear increase in the number of GFP-positive tumor cells was observed three weeks after clone induction (Supplementary Fig. 1 F_1,33_=8.6; P=0.006). Gut dissections confirmed the presence of tumors in the gut (Supplementary Fig. 2). The presence of tumor cells (Apc-Ras clones) had little impact on fly performance and survival over the three weeks of the experimental study (32) (Supplementary Fig. 3).

### Cancer progression and social environment

To investigate the impact of social environment on tumor progression, we exposed adult cancerous females for 21 days, post induction, to various social environments in 40ml food tubes. Individual virgin cancerous females were either kept in tubes alone (social isolation), in groups composed of seven other cancerous flies (homogeneous groups) or in groups with seven non-cancerous control females (heterogeneous groups). Groups of eight control flies were used as a reference (homogeneous group). Tumor growth was significantly affected by the social environment (Wald χ^2^2=6.7, P=0.031). after 21 days we observed that tumor growth was dramatically higher in cancerous flies kept in isolation than in cancerous flies kept in homogeneous groups (Fig. 1). More surprisingly, we also observed that cancerous individual flies kept within a group of control flies showed an increased number of tumor cells compared to cancerous flies grouped together (Fig. 1).

**Figure 1:**
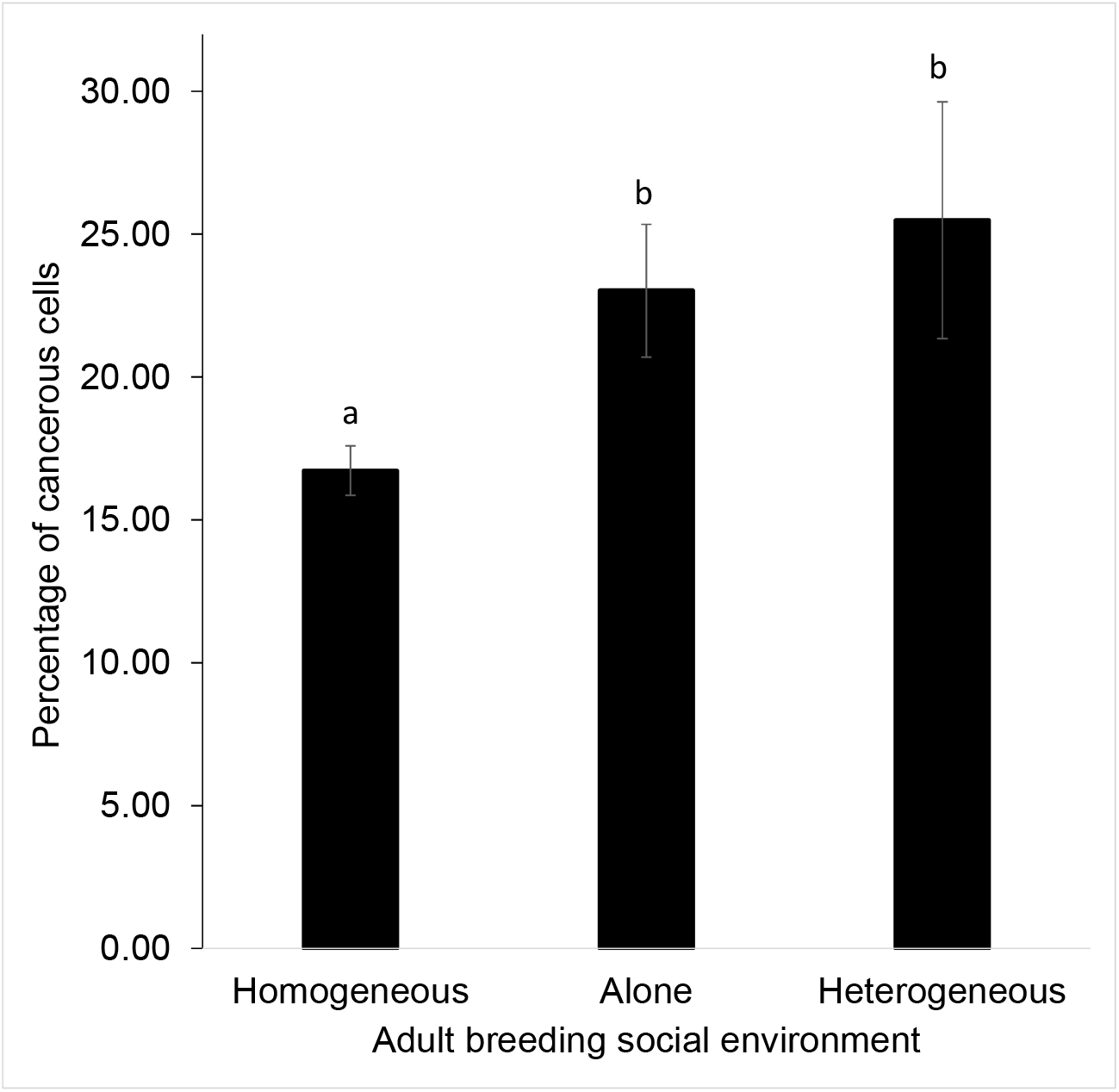
Gut tumor progression as a function of social environment. FACS analysis of GFP-positive cells in guts dissected from 21 days old control or cancerous females as a function of social environment. Error bars: standard error of the mean. N=15 measures for each treatment. Letters are Tukey’s post hoc classification.

### Social interactions

We then analyzed how social interactions were affected by tumor progression and group composition. Using a video tracking setup, we followed the locomotion and interactions of groups of flies (3 weeks post induction) placed in an arena for 1 hour. For social interaction measures we used homogenous groups of eight control or eight cancerous flies, and a heterogeneous group consisting of seven control and one cancerous fly which were kept together for 21 days post induction. Social interaction analyses confirmed that control and cancerous flies had similar locomotor activity independent of their social environment (Fig. 2A; log (trail length). group composition. F_1,84_ = 2.64, P = 0.1; fly state. F_1,84_ = 0.13, P = 0.7; fly state x group composition. F_1,84_ = 3.8, P = 0.061). However, the length of interaction a fly had with another strongly diverged according to group composition and fly state (contact duration. group composition. F_1,84_ = 14.8, P <10^−3^; fly state. F_1,84_ = 26.8, P <10^−3^; fly state x group composition. F_1,84_ = 22.9, P <10^−3^). In homogeneous groups, cancerous flies had longer interactions compared to homogenous control groups (Fig. 2B). Control flies showed the same contact duration whether in homogeneous or heterogeneous groups, whereas cancerous flies showed a strong decrease in contact duration in the presence of control flies (Fig. 2B). Similarly the average number of contact per fly also differed depending on the social context and the state of the flies (number of contact. group composition. F_1,84_ = 17.5, P <10^−3^; fly state. F_1,84_ = 11.4, P = 0.001; fly state x group composition. F_1,84_ = 4.4, P = 0.038). Flies in the control group had the same number of contacts when in the homogenous and heterogeneous groups, while cancerous flies, once again, showed a decrease in the number of contacts when placed with control flies (heterogeneous group) compared to when in a group with other cancerous flies (Fig. 2C). Taken together this would suggest that, in a homogeneous group of cancerous flies, individuals are more aggregated than in a heterogeneous group or a homogeneous group of control flies. We thus concluded that, for a cancerous fly, the composition of the social group strongly affects the level of social interactions. However, our measure of social contact was constrained by the small size of the arena and therefore did not allow us to disentangle the direction of the social contact i.e. which fly showed avoidance and which fly showed attraction.

**Figure 2:**
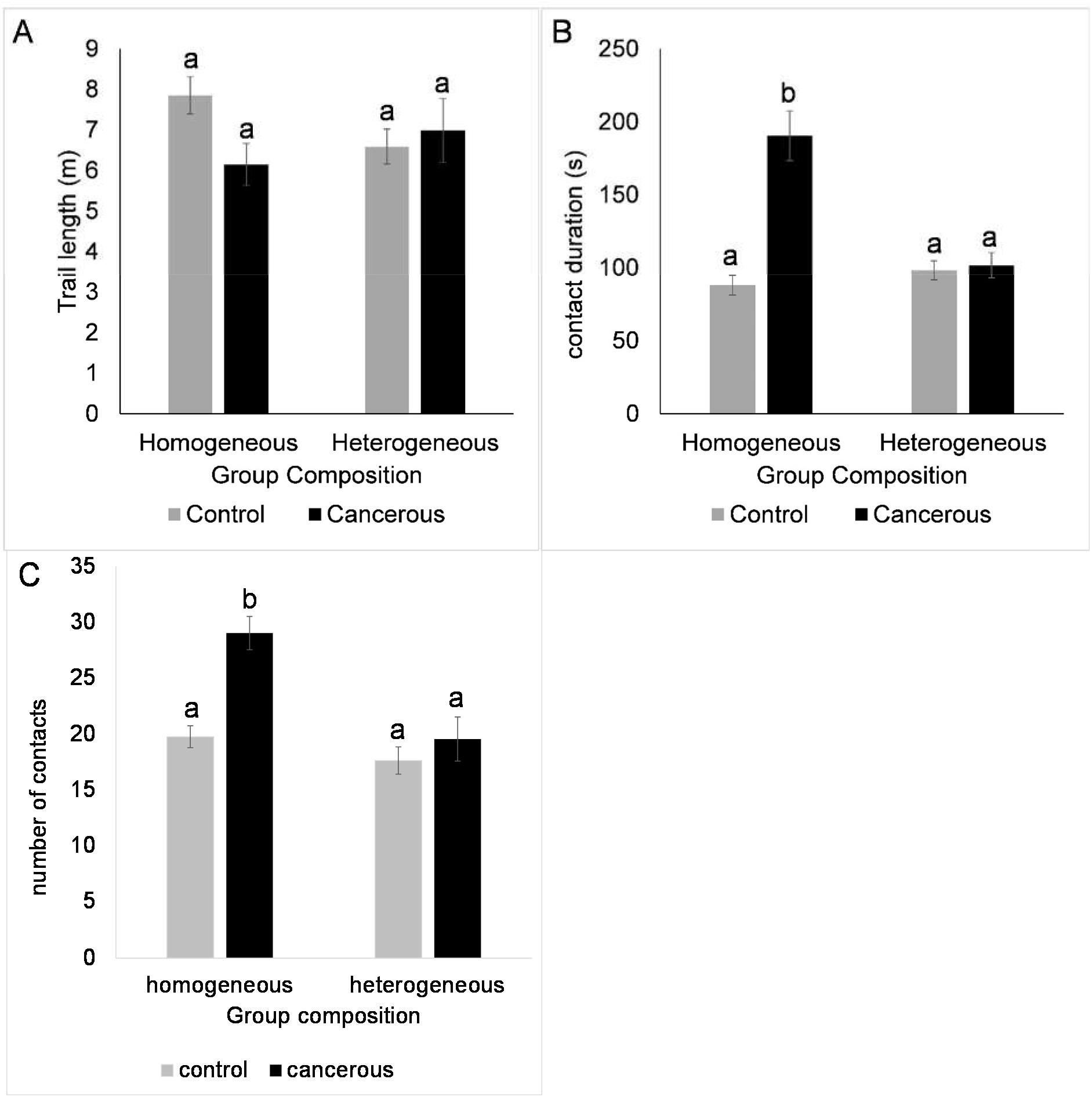
Social interactions for 21 day old females in homogeneous (8 cancerous flies or 8 control flies) or heterogeneous groups (1 cancerous and 7 control flies). 2A: total foraging trail. 2B: averaged cumulated social contact duration one individual has with another individual of the group. 2C: averaged number of contacts one individual has with another individual of the group. Error bars: standard error of the mean. NS: P>0.05; ***: P<10^−3^. N=27 heterogeneous groups and N=18 homogeneous group for each fly state. Letters are Tukey’s post hoc classification

### Cancer progression and social environment choice

Based on the results described above we tested whether cancerous and/or control flies would show variation in social environment choice depending on the level of tumor progression. Using a similar protocol to Saltz (35), we assessed social preference by putting two small mesh cages, each containing 8 “stimulus flies” (cancerous or control) in a plastic, transparent box. The small mesh cages were placed on top of a small petri-dish containing standard food. We introduced a “focal fly” (cancerous or control) in the enclosed box and recorded their position over 7h i.e. whether the fly was found on one of the two mesh cages. Focal and stimulus flies were tested at different ages post heat-shock induction.

Cancerous flies appeared, on average, more attracted than control flies to other cancerous individuals and we observed a general decrease of preference for the cancerous group with age (Fig 3; focal fly: Wald *χ*^2^1=4.1, P=0.04; age: Wald χ^2^1 = 17.6, P<10^−3^; age x focal fly: Wald χ^2^1=2.7, P=0.1).

**Figure 3:**
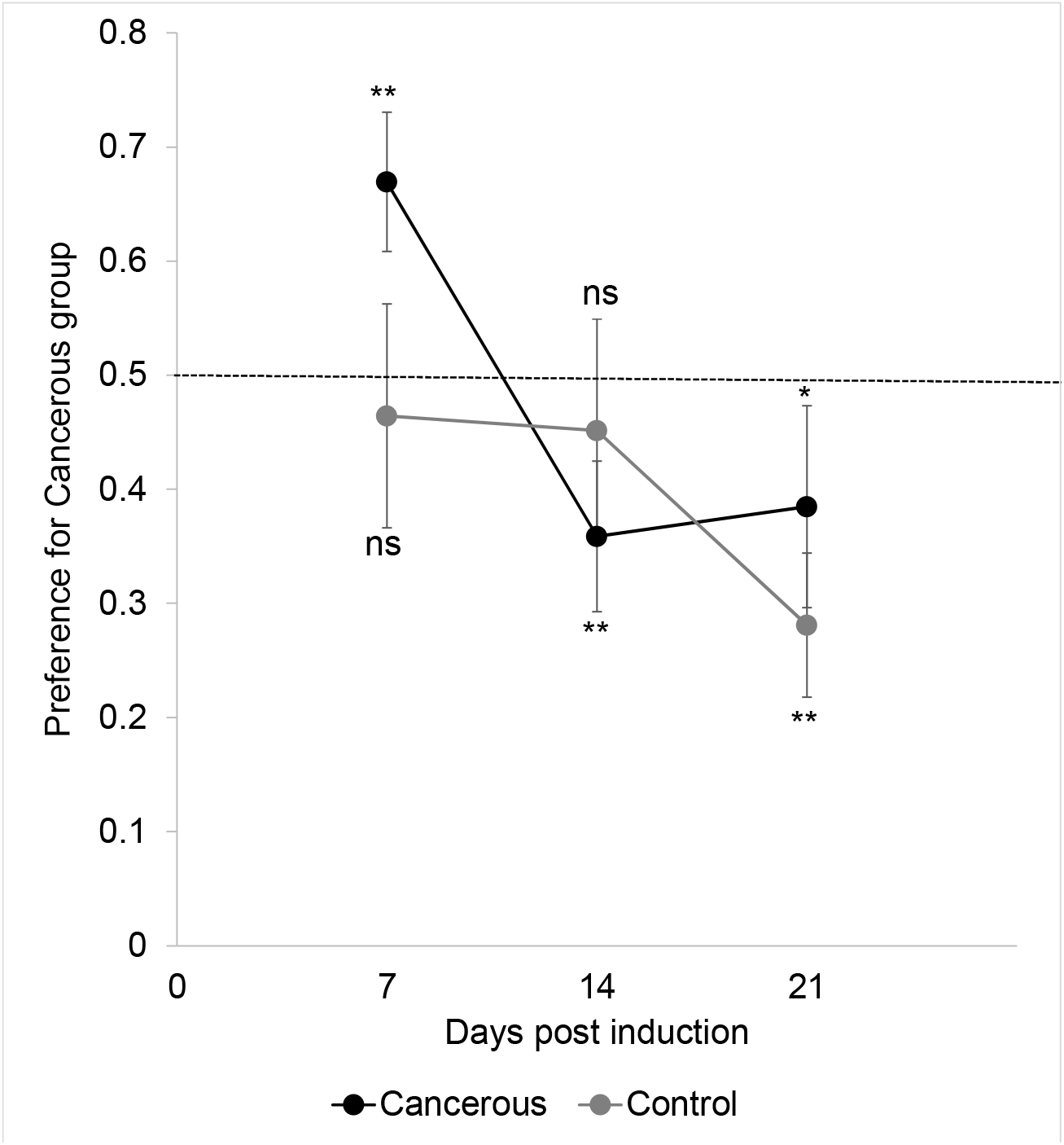
*Dual choice experiment:* proportion of focal flies choosing to land on the mesh cage containing stimulus cancerous flies as a function of age. N=12-21 per treatment. Stars indicate deviation from random choice (binomial test per state and age): ns: P>0.05; *: P<0.05, **: P<0.01; Error bars: standard error of the mean

To understand whether these preferences, in a dual choice, were due to avoidance or attraction, young (7 days post heat shock) focal flies were given a choice between a stimulus group in a mesh cage (8 flies) and an empty mesh cage using a similar experimental design. Cancerous flies showed, on average, attraction for the social group, independent, of the age or the state of the stimulus flies (Fig 4A; intercept: Wald χ^2^1 = 8.1, P<10^−3^; stimulus: Wald χ^2^1=0.06, P=0.79; stimulus age: Wald χ^2^1 = 1.4, P=0.23; stimulus x stimulus age: Wald χ^2^1=0.6, P=0.44). While control flies showed, on average, no clear attraction for the social group, they clearly avoided 3-week-old cancer flies (Fig 4B; intercept: Wald χ^2^1=4.4, P=0.036; stimulus: Wald χ^2^1=2.6, P=0.1; stimulus age: Wald χ^2^1=3.37, P=0.066; stimulus x stimulus age: Wald χ^2^1=6.61, P=0.01).

**Figure 4:**
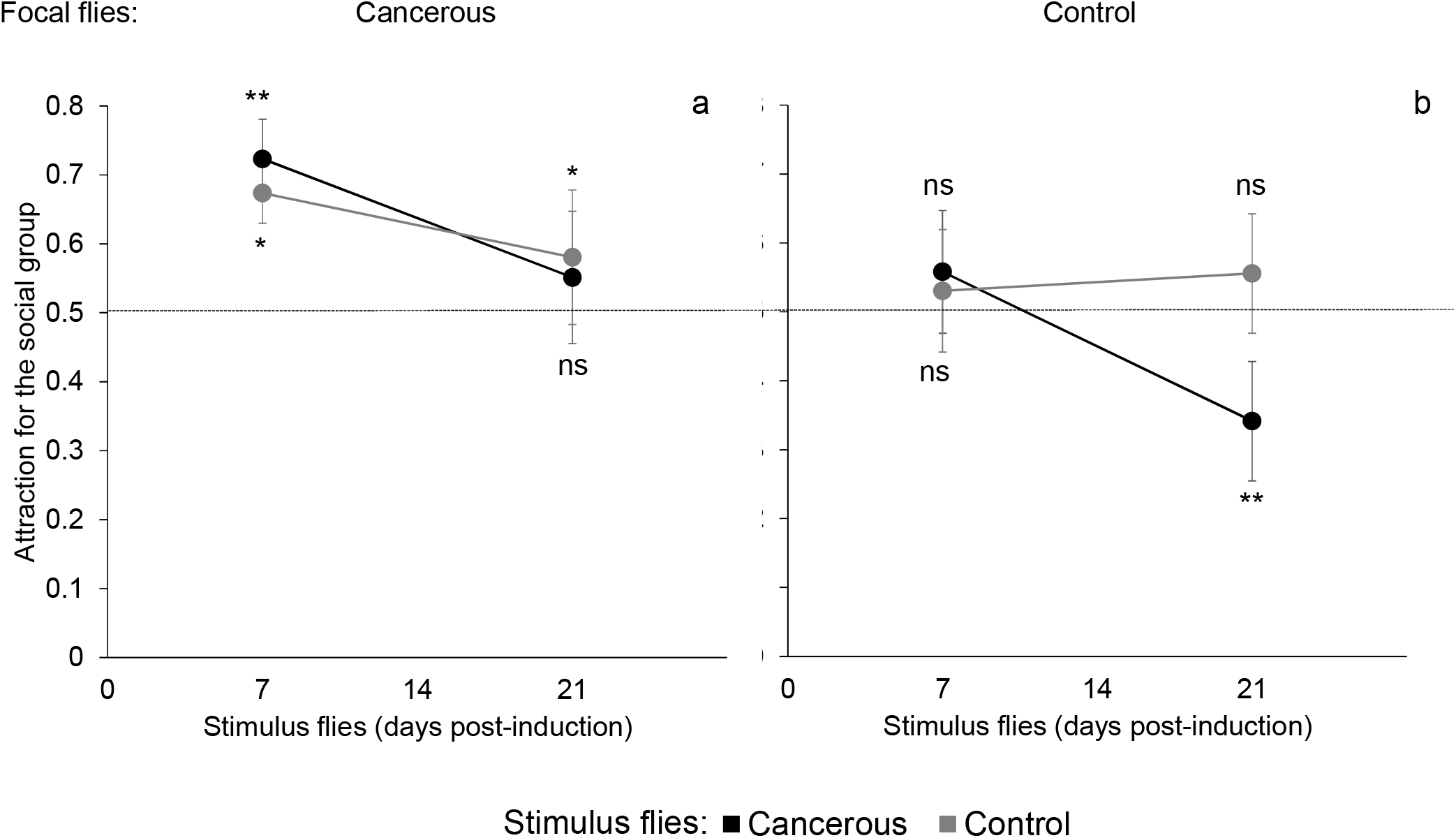
Attraction vs aversion experiment. Proportion of flies (4A: focal cancerous ones; 4B: focal control ones) choosing to land on the mesh cage as a function of age of the stimulus flies. N=16 per treatment. Stars indicate deviation from random choice (binomial test per state and age) ns: P>0.05; *: P<0.05· **: P<0.01.

## Discussion

Here, we show that social environment can significantly shape the development of intestinal-like cancer in Drosophila. Consistent with previous studies on mammals (10, 36), cancerous flies kept in isolation exhibit faster tumor progression than flies kept in groups of other cancerous individuals. However, more importantly we find that variation in group composition also leads to increased proliferation of tumor cells, thus highlighting how subtle variations in social structure may have dramatic effects on the progression of non-transmissible diseases.

Despite the opportunity to interact with others, individual cancerous flies, kept in groups with control flies, developed tumors at similar rates to when cancerous flies were bred in isolation. Social interaction analyses revealed that despite similar locomotor activities, cancerous flies interact considerably less with control flies compared to when they are housed with other cancerous conspecifics. This reduction of social contact may potentially be perceived as a form of social isolation by cancerous flies, which could result in increased tumor growth, analogous to when flies are kept in true isolation. In humans, it has been proposed that the subjective perceived feeling of social isolation may impact psychological and physiological traits as much as real social isolation (37). Furthermore, the social environment choice experiment suggests that control flies may actively recognize and avoid cancerous individuals, especially when tumor growth is significant. Potentially, this may be a result of tumor-induced changes in cuticular hydrocarbons, pheromone profiles or even gut biome. While this avoidance effect may be adaptive for cancer cells, more parsimoniously this behavior could also simply reflect common infection avoidance (38, 39), a behavior that has also recently been observed in Drosophila (40). Even if not contagious, cancerous flies may show particular behaviors, or produce chemical cues, which are generally associated with being sick. While cancerous flies show no such avoidance behavior towards other ‘sick’ flies, there is a decrease in attraction as cancer progresses. We propose that this may be the result of a balance between avoiding other sick individuals and attaining the positive effects of being with other sick individuals (as discussed below).

These findings offer new perspectives on the reciprocal relationship between disease and social behavior. While we observe that social structure has profound effects on disease progression, our study also suggests that disease might play a fundamental role in influencing group composition. We found that cancer developed at a slower rate when flies were with other cancerous flies. Moreover, we observed that cancerous flies, instead of showing avoidance behavior towards other cancerous flies as non-cancerous flies did, exhibit strong social attraction towards each other, especially at the beginning of tumor development. This raises questions on the very early impact of internal oncogenic process on individual behavior and natural selection pressures on this process (41). While, at this stage, tumors are not found to affect the survival of flies, previous research suggests that cancer may affect fitness in other ways. Female Drosophila, bearing colorectal tumors, bring forward their peak oviposition period suggesting that flies are adapted to minimize the costs of cancer on fitness (18). The social behavior of cancerous flies found in this study could also be interpreted as an adaptive process, which reduces cancer progression. Further studies would be necessary to determine the exact proximate factors responsible for the effect found in this study, as well as the extent to which generalizations can be made across other cancer types (or indeed other illnesses) and animal species.

Our findings highlight the importance of social structure on disease progression, beyond the context of transmission. This is the first time that a direct link between social environment, specifically group composition, and cancer progression has been shown, while removing all other confounding psycho-sociological parameters that are frequently encountered in human studies. More generally, this study brings new light to how sociality impacts health and fitness in animals and its potential implication in human disease therapy. Moreover, we provide essential data to the emerging topic of evolutionary ecology of cancer, and demonstrate the importance of neoplasia as a fitness limiting factor that potentially influences life history adaptations and strategies.

## Author Contributions

E.H.D, T.B, C.M., FT, J.M and F.M designed the experiments. E.H.D., T.B., J.D.S., C.M., M.S. and J.M. performed the experiments. C.S, J.M and F.M analyzed the data. E.H.D., J.M., F.T., A.C., C.S., B.U. and F.M. wrote the manuscript.

## Acknowledgements

This work was supported by the ANR (Blanc project EVOCAN to FT and project DROSONET to FM and CS), the CNRS (INEE and INSB), *Fondation ARC* (1555286 to JM and FM), The French league against Cancer (M27218 to JM), IDEEV program (to FM), by an International Associated Laboratory Project France/Australia, by the French-Australian Science Innovation Collaboration Program Early Career Fellowship (BU), by André Hoffmann (Fondation MAVA), Fyssen Foundation (to FM and EHD) and the French Government (fellowship 2015-155 to MD). We thank F. Bastin, J.C. Sandoz and Couto for their help with confocal imaging.

## Methods

### Drosophila stocks and genetics

*yw,HS-flp;esg-gal4,UAS-GFP;FRT82B,Tub-Gal80* (line 1), *yw,HS-flp;UAS-Ras^V12^;FRT82B,Apc2^M175K^,Apc^Q8^* (line 2) and *yw,HS-flp;;FRT82B* (line 3) flies (32) were balanced over co-segregating *SM5-TM6B* balancers. In all experiments, cancerous flies were *HS-flp;esg-gal4, UAS-GFP/UAS-Ras^V12^;FRT82B,Tub-Gal80/FRT82BApc2^N175K^,Apc^Q8^* (offspring 1 of line 1 crossed to line 2), whereas controls were *HS-flp;esg-gal4,UAS-GFP;FRT82B,Tub-Gal80/FRT82B* (offspring 2 of line 1 crossed to line 3). In this study, MARCM clones (32) were randomly generated in heterozygous flies by flipase-induced exchange of pairing chromosome arms, resulting in mosaic individuals where homozygous *Apc2^N175K^, Apc^Q8^* mutant cells lacked the Gal80 repressor, thereby allowing Gal4 activity and the subsequent expression of GFP and Ras^V12^ for clones located in intestinal progenitor cells. Conversely, MARCM control clones are wild type for both Apc genes and do not express Ras^V12^. MARCM clones were generated by a 1 hr heat shock at 37°C of 3 days old females (32). Several attempts were made to use non-induced (no heat shock) offspring flies as controls, however, due to unpredictable tumor appearance (a few of flies developing tumor without heat shock) the lineage was declared not suitable as a reliable control.

### Flow cytometry

Prior to the quantification of GFP-positive cells as an estimate of tumor progression, flies were starved overnight, provided only with water. The entire midgut and the Malpighian tubules, that also exhibit tumor-like structures, were dissected in PBS (Phosphate buffer saline). Samples of five guts, each taken from a separate tube, (for example, one replicate of the homogenous treatment consisted of five guts of cancerous flies, each of these five flies randomly taken from five different homogeneous tubes) were digested by collagenase (125µg in 60µl PBS) (Sigma-Aldrich) for 2h at 27°C with gentle agitation. Samples were then complemented with 10 µl Trypsin 10X (Sigma-Aldrich) and nuclei were stained by Hoechst 33342 (0.5µg/ml) for at least 1hr. Samples were filtered and analyzed on a Partec PAS III (Figure S1).

### Social environment

flies were sexed at emergence and control or cancerous females were kept in groups until the third day post emergence. Control and cancerous virgin females were heat shocked at 37°C for 1h. Flies were then introduced into new 40ml food tubes according to their treatment. Control and cancerous flies were partially wing-clipped on the right or left wing to distinguish their genotype. Previous behavioral studies have shown that wing clipping has no effect on social interactions (42). Flies were then kept at 25°C on standard food (changed every 3 days) until day 21 post induction. Note that the size of the tubes was small (40ml) enough to limit any effect of complete social isolation. Tumor size was estimated with flow cytometry. Data were analyzed with generalized linear model (binomial distribution, Pearson correction for over-dispersion) on the number of tumor cells vs the total number of cells counted. Tukey’s post hoc tests were then applied.

### Social interactions

Each group (composed of flies taken from different food tubes i.e. had never previously interacted together) was introduced into a semi opaque white polyoxymethylene (Delrin) arena (diameter 100mm; height 5mm) covered with a transparent Plexiglas for 1h. Our experimental design allowed us to simultaneously track 4 groups of 8 flies over the 4h. The tracking apparatus consisted of four synchronized firewire cameras (Guppy pro, Allied vision technologies), each filming one interaction arena that was backlit by a 150X150 mm IR backlight (R&D vision). We used Vision software to analyze spatial data (open-source C-trax 0.3.7 (43)) that allowed us to collect 10 positions per second for each fly over 1h video experiments. Tracking corrections were made post C-trax analysis with fixerrors Matlab toolbox (Ctrax-allmatlab version 0.2.11) using Matlab software 7.11.0 to suppress swaps between individuals. We then calculated, for each fly, the total length of the path, the distance to other flies, the number of contacts with other flies (a contact was considered when the distance between the centers of two individuals was smaller than, or equal to, one mean body length of the individuals for 1s or more) and the duration of each contact. These values were averaged to obtain, for each replicate group, one value for cancerous state and/or control state. For each measure we performed a general linear model including as fixed explanatory variables group composition (homogeneous vs heterogeneous), fly state (cancerous or control) and the interaction group composition x fly state. Tukey Post-hoc contrasts among treatments were tested.

### Social environment choice

flies were sexed at emergence and control or cancerous females were kept in groups until the third day post emergence. Control and cancerous virgin females were heat shocked at 37°C during 1h and kept in group until the day of the experiment. The experimental setup consisted of a 17x12x5cm plastic box in which 2 small 2x2x2cm mesh cages were introduced and each placed on a 3cm diameter petri dish containing standard food. The two cages were positioned at opposite ends of the box. Groups of eight flies were placed in the mesh cages. In the dual choice experiment, one mesh cages contained control flies whereas the other contained cancerous flies. In the attraction vs avoidance experiment only one of the two cages contained stimulus flies. A focal fly (control or cancerous) was introduced in the box 15h before starting the experiment. The position of the fly was then visually recorded every 30min between 10am and 5pm. We only considered cases where the fly was positioned on a mesh cage or the associated petri dish. For the dual choice experiment, focal and stimulus flies were of the same age (7, 14 or 21 days post induction), whereas for the attraction vs aversion experiment the focal fly was always 7 days old post induction and the stimulus flies were either 7 or 21 days post induction. The number of times a focal fly was observed on a cancerous stimulus cage (for the dual choice experiment) or the stimulus cage (for the attraction vs aversion experiment) compared to the total number of cage landings were then analyzed with a general linear model and a binary logistic regression. For the dual choice experiment, we first compared the behavior of cancerous vs control flies: State of the focal fly was included as a fixed factor and fly age was included as a covariate. For the attraction vs aversion experiment we separately analyzed the behavior of each focal fly (i.e. cancerous or control) as a function of stimulus fly state and age. Finally, a binomial test, for each independent measure, was performed to test for a significant deviation from random choice.

## SUPPLEMENTARY INFORMATION

**Figure SI:**
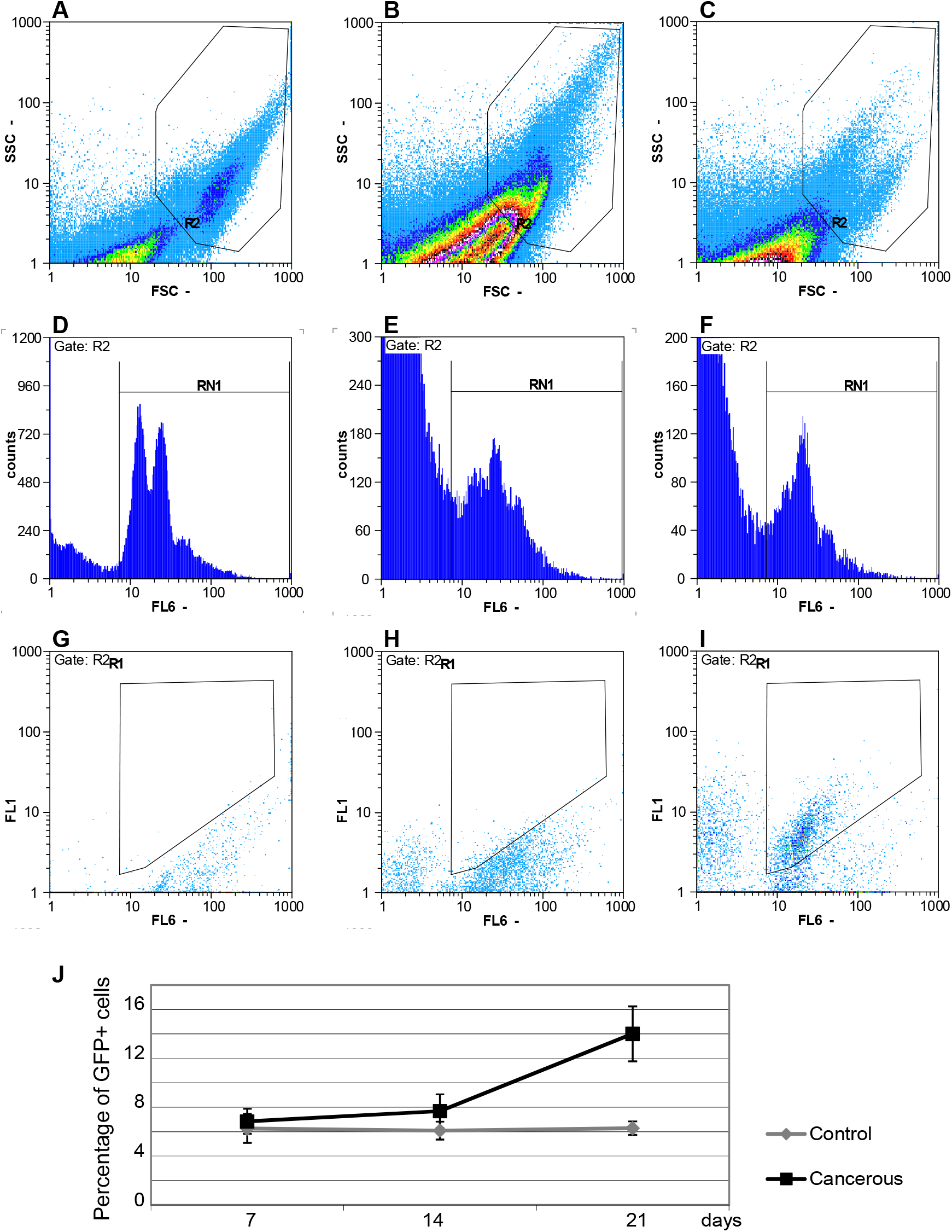
FACS analysis of intestinal cells. (A-C) SSC versus FSC Dotplots to gate (R2) the cells of interest from dissected imaginal discs (A) or guts of *w^1118^* (B) and cancerous (C) flies. (D-F) Selection of the *Drosophila* cells within the R2 gate, with respect to their DNA content (RN1); imaginal disc cells allow to identify diploid G1 and G2 cells (2 major pics in D); the intestinal cells of *w^1118^* (E) and cancerous (F) flies, which undergo DNA endoreplication contain at least the DNA content of G1 imaginal disc cells. (G-I) FL6 (DNA) versus FL1 (GFP) Dotplots to select the GFP-positive cells from gut of cancerous flies (R1 in I), whereas the R1 gate is empty when analyzing imaginal disc cells (G) and gut cells (H) of *w^1118^* flies. (J) FACS quantification of GFP-positive cells in cancerous (black line) and control (grey line) guts dissected from adult females at 7, 14 and 21 days past heat shock-induced clonal recombination. Genotype of cancerous and control flies are *HS-flp;esg-gal4,UAS-GFP/UAS-Ras^V12^;FRT82B,Tub-Gal80/FRT82BApc2^M175K^,Apc^Q8^* and *HS-flp;esg-gal4,UAS-GFP;FRT82B,Tub-Gal80/FRT82B*, respectively. Error bars: standard error of the mean. N=8 measures for each treatment.

**Figure S2:**
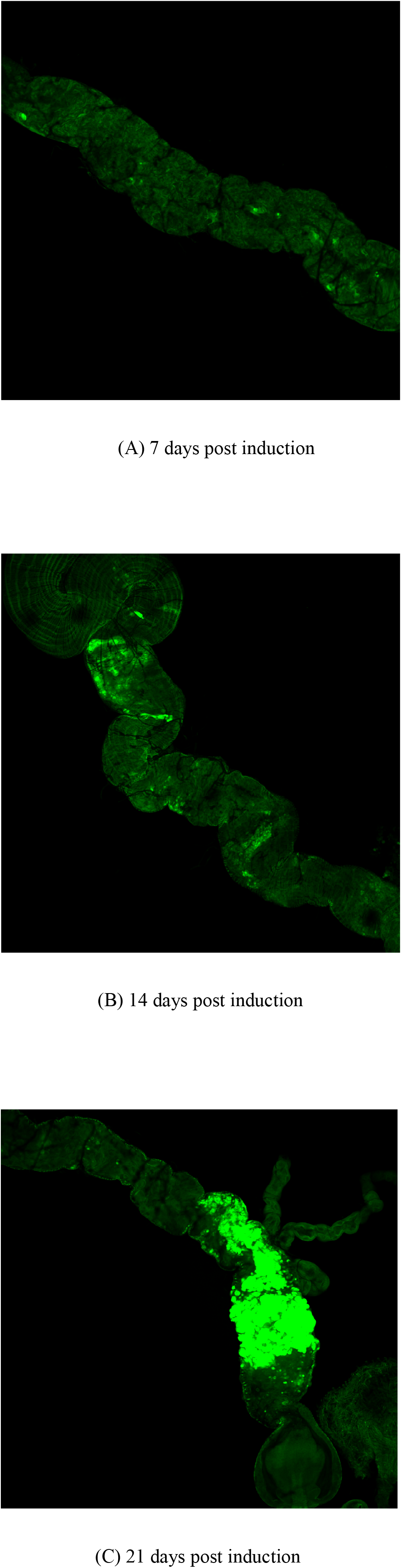
Progression of tumor growth in the Midgut. (A-C) APC-Ras clones in the midgut labeled by GFP (green) at 7 days (A), 14 days (B) and 21 days (C) after clonal-induced recombination. Note that 7-day old clones mostly appear as isolated cells, whereas 14-day old clones appear as groups of few cells; 21-day old clones invade large portions of the midgut. Guts from overnight starved flies were dissected in PBS and fixed with 3,7% formaldehyde in PBT (PBS, 0,1% Tween 20) for 20 mn at room temperature; guts were then extensively washed and mounted in DABCO (sigma). Image acquisitions were obtained using laser scanning confocal microscope (LSM700; Carl Zeiss, Jena, Germany) and a solid-state 488-nm laser for exciting GFP. Image were acquired at a resolution of 1024x1024 pixels using a water immersion objective (x10 achroplan 0.3 NA). **Fly physical performance:** We compared physical and behavioral performances between control and cancerous flies at different ages by exposing them to a ‘negative geotaxis’ behavior test. This involved repeatedly tapping a tube of flies, which causes them to fall, and then measuring the number of individuals that show an escape response (ascending the walls of the tube) over time. When repeatedly performed, this escape response tends to diminish due to exhaustion. This test has been extensively used as a robust proxy to estimate physical and behavioral performance and locomotor activity. We followed a protocol based on the one initially developed by JW Gargano and co-workers (44) and modified by MJ Tinkerhess (45). Groups of 10 cancerous or control females (7, 14 or 21 days post induction) were introduced into 40ml tubes and placed vertically on a platform which automatically taps the tubes every 4 seconds causing all flies to fall to the bottom of the tube. Every 15 min we visually recorded the number of flies in each tube showing negative geotaxis (climbing at least 2/3 of the tube between two taps). The experiment lasted 2 hours. The data were analyzed with repeated measures ANOVA including time (repeated measure), fly state (cancerous or control) and age (covariate 7,14 or 21 days post induction) as factors. The presence of tumor cells (Apc-Ras clones) has little impact on fly performance and survival over the three weeks of the experimental study. Cancerous and control flies subjected to repeated taps over two hours showed a progressive decline in negative geotaxis response (Repeated measure ANOVA: time: F_1,50_ = 319,9 P<10^−3^; Fig. S3) which was stronger as flies got older (age: F_1,50_ = 63.1 P<10^−3^; time x age: F_1,50_ = 8.8; P = 0.004). However, no difference between cancerous or control flies could be observed at any age post-induction (state: F_1,50_ = 0.73; P = 0.39; state x age: F_1,50_ = 0.17; P = 0.68) suggesting that despite the general observation of age related impairments, Apc-Ras-induced tumors do not affect physical performance and locomotion 3 weeks after induction.

**Figure S3:**
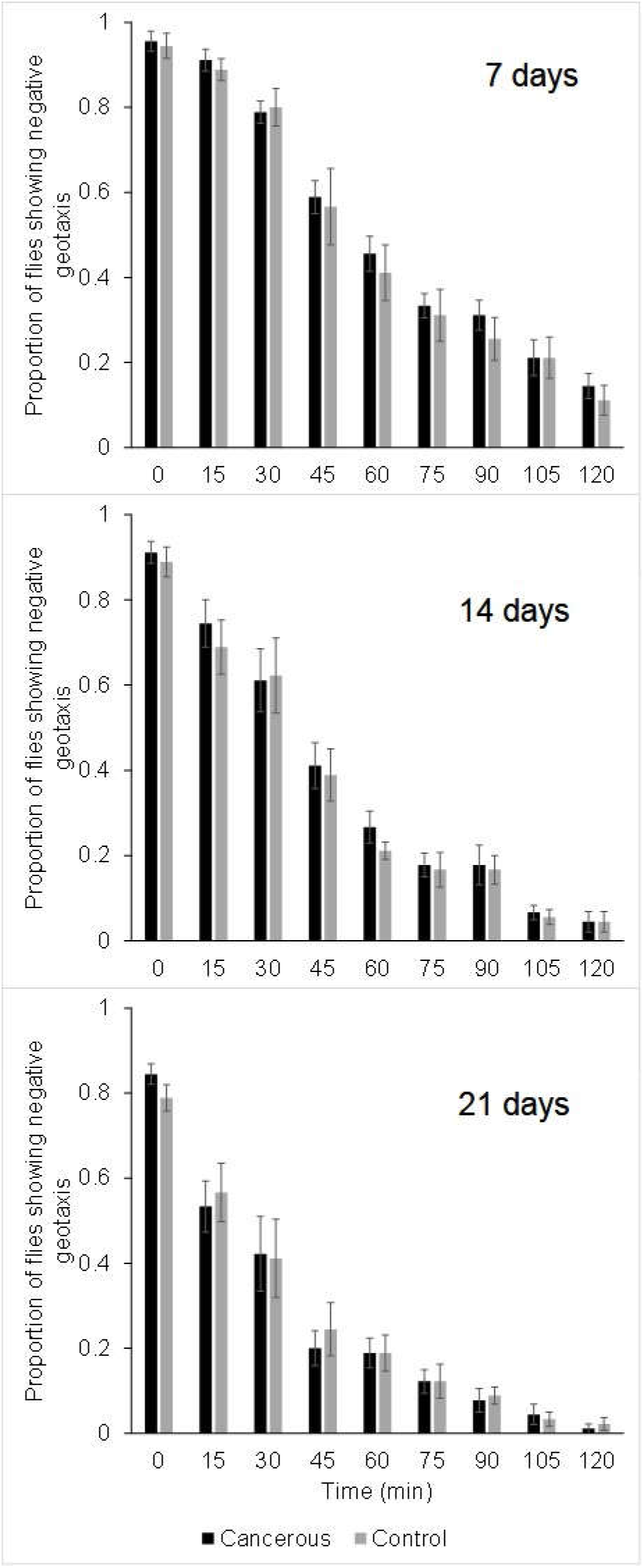
Repeated ‘negative geotaxis’ behaviour test performed at different ages for cancerous and control flies. Proportion of flies ascending the walls of the tube after it has been repeatedly tapped. Measures were taken every 15min over 2h at different ages (7, 14 or 21 days post induction). Error bars: standard error of the mean. N=18 groups followed during 2h for each treatment and age

